# Construction and Elaboration of Autobiographical Memories from Multiple Visual Perspectives

**DOI:** 10.1101/317594

**Authors:** Heather Iriye, Peggy L. St. Jacques

## Abstract

Visual perspective, recalling events from one’s own eyes or from an observer-like viewpoint, is a fundamental aspect of autobiographical memory (AM). Yet, how visual perspective influences the functional mechanisms supporting retrieval is unclear. Here, we used a multivariate neuroimaging analysis to characterize the spatiotemporal dynamics supporting AM retrieval from multiple visual perspectives. Both own eyes and observer perspectives engaged an AM retrieval network (i.e., hippocampus, anterior and posterior midline, lateral frontal and posterior cortices) that peaked during later retrieval periods but was recruited less strongly for observer perspectives. Functional connectivity analyses with an anterior hippocampal seed revealed that visual perspective also altered interactions among neural regions and their timing during retrieval. There was stronger hippocampal connectivity with a posterior medial network during the initial construction of AMs from observer perspectives and stronger connectivity with a medial temporal lobe network during later retrieval periods from own eyes perspectives, suggesting that visual perspective directs how neocortical systems guide retrieval. Our findings demonstrate that visual perspective influences AM retrieval by altering hippocampal-neocortical interactions and subsequently the strength of neural recruitment in the AM retrieval network during later retrieval periods, thereby supporting the central role of visual perspective in shaping the personal past.

---

Autobiographical memory (AM) retrieval requires taking a particular egocentric perspective, or window on which to remember the past. AMs can be retrieved from the visual perspective of one’s own eyes (OE) or from multiple observer (OB) vantage points, and the particular visual perspective adopted shapes the content and phenomenology of memories (e.g., Berntsen and Rubin 2006; Butler and others 2016; Marcotti and St Jacques 2018; McIsaac and Eich 2002; Nigro and Neisser 1983; Rice and Rubin 2009). Previous functional neuroimaging studies have indicated that AM retrieval is involves neural recruitment in anterior and posterior midline regions, medial temporal lobe (MTL), and frontoparietal regions, which overlaps with the default network (Andrews-Hanna and others 2014; Kim 2012; St Jacques and others 2011b). Moreover, this research has shown that the pattern of functional recruitment and functional interactions within these regions varies across early and late phases of AM retrieval (Addis and others 2007; Daselaar and others 2008; Inman and others 2018; McCormick and others 2015; St Jacques and others 2011b). A significant gap in our understanding, however, is that the majority of studies of AM either do not manipulate visual perspective or have focused solely on memories retrieved from an OE perspective. Thus, it is currently unknown whether visual perspective shapes memories by modifying how a specific memory is initially searched for and constructed or by biasing the subsequent elaboration of details within memories. Here, we take a multivariate fMRI analysis approach to investigate the influence of visual perspective on functional neural recruitment and functional connectivity across the time-course of AM retrieval.

A growing number of neuroimaging studies have begun to investigate visual perspective during long-term memory retrieval (Freton and others 2014; Grol and others 2017; Hebscher and others 2018; St Jacques and others 2018; St Jacques and others 2013; St Jacques and others 2017). Structural neuroimaging studies have shown a positive relationship between precuneus volume and the tendency to adopt an OE perspective during AM retrieval (Freton and others 2014; Hebscher and others 2018). However, functional neuroimaging studies of memory retrieval are less consistent—with some studies finding similar involvement for OE and OB perspectives (Eich and others 2009; St Jacques and others 2018) and others finding greater involvement for OB than OE perspectives (Grol and others 2017). One reason for this inconsistency may be that the precuneus is involved in both the representation and manipulation of mental images that support the adoption of OE and OB perspectives alike (for reviews see Byrne and others 2007; Cavanna and Trimble 2006). Supporting this viewpoint, St. Jacques and colleagues (2017) demonstrated that the precuneus contributes to the ability to shift to alternative visual perspectives—irrespective of the direction of perspective shifting (St Jacques and others 2018). Visual perspective also influences neural recruitment in regions associated with bodily self-representation and emotional response, consistent with reports of heightened emotional intensity and physical sensations that accompany retrieval from an OE perspective (e.g., Berntsen and Rubin 2006; McIsaac and Eich 2002). For example, Eich and colleagues (2009) asked participants to retrieve memories for complex lab-based events and found differential involvement of amygdala, somatomotor, and insular cortices for memories retrieved from OE compared to OB perspectives, which they linked to behavioral decreases in affect and physical sensations during retrieval from an OB perspective. Visual perspective may also influence functional recruitment of the hippocampus and the integrity of its network connections. Retrieving memories encoded from OE compared to OB perspectives alters neural recruitment of the hippocampus during remembering (Bergouignan and others 2014), and higher OE ratings during AM retrieval are associated with greater engagement of an MTL network centered on the hippocampus (St Jacques and others 2013). However, previous research has not delineated how explicitly manipulating visual perspective on a trial-by-trial basis influences the time course of AM retrieval.

Construction and elaboration of AMs involve dissociable neural mechanisms (Addis and others 2007; Daselaar and others 2008; Inman and others 2018; St Jacques and others 2011b), which may be differentially affected by visual perspective. On the one hand, visual perspective may influence AM retrieval during the initial construction of memories, because perspective is tightly linked to self-referential processes associated with the medial prefrontal cortex (PFC; e.g., D’Argembeau 2013) that emerge early during the time course of event retrieval and initiate neural recruitment in a widespread network, including the hippocampus (McCormick and others 2015; St Jacques and others 2011b). Additionally, manipulating visual perspective during AM retrieval involves similar constructive demands as those that support episodic simulation (St Jacques and others 2018), which rely on frontoparietal regions (for metaanlysis see Benoit and Schacter 2015), and these constructive processes tend to emerge earlier (Addis and others 2007). Finally, emotional intensity, which is influenced by visual perspective during AM retrieval (e.g., Berntsen and Rubin 2006), modulates neural recruitment in the amygdala and hippocampus during memory construction (Daselaar and others 2008). On the other hand, visual perspective may predominately influence later elaboration of AMs, when sensory and perceptual aspects of memories are re-experienced (Conway and Pleydell-Pearce 2000). Visual perspective is supported by mental imagery processes linked to the precuneus and visual cortices (Cavanna and Trimble 2006), that emerge during the later elaboration of memories (Daselaar and others 2008; McCormick and others 2015). Moreover, during elaboration there is also heightened functional connectivity between the precuneus and posterior visual cortices (Inman and others 2018), and bilateral posterior hippocampus and visual cortices (McCormick and others 2015). Alternatively, visual perspective could influence both phases of AM retrieval. For example, St. Jacques and colleagues (2013) showed that people who had higher OE ratings during AM retrieval recruited the MTL network more during both construction and elaboration phases. While several fMRI studies have investigated differences between construction and elaboration of AMs during retrieval (e.g., Daselaar and others 2008; McCormick and others 2015; St Jacques and others 2011b), when precisely visual perspective affects retrieval has yet to be determined. Thus, it is unknown whether visual perspective influences how AMs are initially selected or, alternatively, how specific memory details are elaborated upon during later phases of retrieval.

The present fMRI study examined how explicitly manipulating visual perspective during AM retrieval influences the neural mechanisms associated with construction and elaboration of memories. Participants retrieved AMs from a specified OE or OB perspective elicited by familiar location cues, or completed a control condition involving spatial visualization without retrieving a specific AM. A multivariate analytical approach, partial least squares (PLS; McIntosh and Lobaugh 2004), was employed to examine task-related differences in patterns of brain activity. Additionally, we conducted a PLS functional connectivity analysis on a seed region in the hippocampus because functional integration with this region has been shown to differ across construction and elaboration phases of AM retrieval (McCormick and others 2015) and is also influenced by visual perspective (St Jacques and others 2013). Critically, the use of PLS also enabled us to distinguish task-related differences at specific time points during AM retrieval in order to distinguish early (i.e., construction) from later (i.e., elaboration) AM retrieval periods. Based on the link between visual perspective and mental imagery processes that occur during elaboration of AMs (Daselaar and others 2008), we predicted that visual perspective would primarily influence later periods of AM retrieval, as reflected by greater differences in the pattern of neural activity for AMs retrieved from OE and OB perspectives during later time periods. Additionally, given evidence that visual perspective influences engagement of MTL networks that contribute to AM retrieval across both construction and elaboration phases (St Jacques and others 2013), we predicted that visual perspective would influence hippocampal-connectivity across both early and later stages of retrieval.

## Materials and Methods

### Participants

Participants included 25 healthy, right-handed young adults (age range: 18 to 30 years) with no prior history of neurological or psychiatric impairment, and who were not currently taking medication that affected mood or cognitive function. Participants provided informed written consent as approved by the School of Psychology at the University of Sussex. Five participants were excluded from the analysis due to issues during the fMRI session (i.e. movement greater than 3 mm, n = 3; not following instructions, n = 1; and technical issues during data collection, n = 1). Thus, the final analysis was performed on 20 participants (8 women; mean age in years = 22.5, *SD* = 2.89).

### Procedure

The study involved two separate sessions. In a pre-scanning session, participants were asked to provide titles for 130 distinct and sufficiently specific spatial locations they had visited in the last three years (e.g., *arcade of the Brighton Pier* versus Brighton Pier). Each location was rated according to familiarity, vividness, and emotional intensity on 7-point scales from 1 = low to 7 = high, as well as the date of the last visit (1 = > 2 years, 2 = last 2 years, 3 = last year, 4 = last 6 months, 5 = last month, 6 = last week, 7 = today). The spatial locations were then randomly assigned to one of six experimental conditions (described below) and matched across the subjective ratings. Thus, there were no initial differences in the nature of the spatial location cues within each condition (see Supplementary Table 1).

The fMRI scanning session took place five to nine days later (mean in days = 6.63, *SD* = .96). On each trial, participants were presented with a spatial location cue and asked either to retrieve a specific AM that had occurred at that location (AM task) or to mentally visualize the spatial location without retrieving a specific AM (control task). To distinguish between construction and elaboration phases, participants were asked to press a button once they had thought of a specific AM or could visualize the spatial location, and then to continue to elaborate upon the memory or spatial visualization in as much detail as possible. Participants were given 17.5 s for the entire retrieval or visualization period.

To investigate the influence of visual perspective on construction and elaboration phases of AM retrieval, we asked participants to adopt OE or OB perspectives. We also manipulated the typicality of the visual perspective in order to better understand how adopting multiple visual perspectives influences AM retrieval (Rice and Rubin 2011). In the OE perspective conditions, participants were asked to retrieve AMs as if they were seeing the event from their own eyes, either from a typical OE vantage point or from an atypical OE vantage point in which they mentally rotated the scene by switching left and right. In the OB conditions, participants were asked to retrieve AMs as if seeing themselves in the event, outside of their body at a distance within six feet, either from a typical OB vantage point at eye level or from an atypical OB vantage point at floor level. In the control task, participants were also asked to visualize the spatial location from a typical vantage point that included the proximal aspects of the location (e.g., zooming in on the location within the Brighton Pier), or from an atypical vantage point that included the distal aspects of the location (e.g., zooming out on the location of the Brighton Pier in relation to the city of Brighton).

Immediately following each trial, participants were asked to provide subjective ratings of vividness, emotional intensity, and perspective maintenance (i.e., the ease with which the specified perspective was sustained), each on 4-point scales from 1 = low to 4 = high. Participants had 2.5 s for each rating and responded using a four button MRI-compatible response box. Prior to scanning, participants conducted a practice session to familiarize them with the study conditions and timings of the responses.

There were six functional runs consisting of either 18 trials (four runs) or 24 trials (two runs), for a total of 20 trials per condition. Trial order was pseudo-randomized, such that no condition was repeated more than twice consecutively. Trials were separated by an active baseline consisting of a left versus right decision task, which was equally spaced across a variable length (2.5 s to 10 s; Stark and Squire 2001) and distributed exponentially such that shorter inter-trial intervals occurred more frequently than longer.

### MRI Data Acquisition and Preprocessing

Functional and structural images were collected on a 1.5 Siemens MRI scanner. Detailed anatomical data were collected using a multi-planar rapidly acquired gradient echo (MPRAGE) sequence. Functional images were acquired using a T2*-weighted echo planar sequence (TR = 2500ms, TE = 43ms, FOV = 192 mm × 192 mm). Whole-brain coverage was obtained via 33 coronal oblique slices, acquired at an angle corresponding to AC-PC alignment in an ascending fashion, with a 3 × 3 mm in-plane resolution.

Preprocessing of functional images was performed using SPM12 (Wellcome Department of Imaging Neuroscience, London, UK) using standard methods. Functional images were corrected for differences in acquisition time between slices for each whole brain volume using slice-timing, realigned within and across runs to correct for head movement, spatially normalized to the Montreal Neurological Institute (MNI) template (resampled at 2 × 2 × 2 mm voxels), and then spatially smoothed using a Gaussian kernel (8 mm full-width at half maximum).

### Behavioral Data Analysis

We conducted a 3 (Condition: OE, OB, Spatial) × 2 (Visual Perspective: Typical, Atypical) repeated measure ANOVA separately on reaction time to construct the event and online subjective ratings (i.e., vividness, emotional intensity, and perspective maintenance). Post-hoc comparisons were Bonferroni corrected and appropriate tests were applied when assumptions of sphericity were violated. One participant was excluded from the behavioral analyses of the subjective ratings due to an insufficient number of responses.

### fMRI Analysis: Partial Least Squares

To analyze the fMRI data, we used spatiotemporal PLS, a multivariate analysis approach that determines the optimal least squares fit between task conditions and distributed brain activity during experimental trials (McIntosh and Lobaugh 2004). The advantage of using PLS in the current study is that it characterizes changes in task-related brain activity at multiple time lags across the length of the experimental trial. Here, we specified a 17.5 s temporal window (i.e., 7 time lags, each equal to 2.5 s or 1 TR), to examine the multivariate pattern of brain activity during the retrieval or visualization period.

A cross-covariance matrix was created for each trial, including all participants and conditions, based on a design matrix containing condition information and a data matrix containing brain voxels. Singular value decomposition was applied to the cross-covariance matrix resulting in a set of extracted orthogonal components or latent variables (LVs) that provide the best fit of the data. Each LV reflects a set of contrasts that characterize the differences or similarities among the task conditions, and an associated pattern of distributed brain activity by time lag. LVs are associated with singular values, or the proportion that each LV contributes to the overall covariance between task conditions and brain activation. Each voxel is linked to a salience score, dependent on the observed covariances. Each salience can then be multiplied by the blood oxygen level dependent (BOLD) signal value of its associated voxel and summed across all voxels to yield a brain score, which can then be used to compare patterns of brain activity across the task conditions. Greater activity in brain regions with positive saliences on a LV will produce positive mean brain scores for a given condition over each time point, while negative saliences will produce negative mean brain scores. LVs are progressively extracted in successively smaller amounts until all the data are accounted for.

The statistical significance of the LVs was determined using 500 permutation tests (e.g., Addis and others 2004; Burianova and others 2010), which applies singular value decomposition to calculate a new set of LVs after randomly re-ordering the data matrix rows. The newly obtained singular value of each LV is then compared to the originally derived value to create an updated weighted probability of the original singular value, dependent on the number of times it is exceeded by the results of each permutation test (McIntosh and others 1996). The reliability of the results was assessed by 300 bootstrap estimations on the standard error of the saliences for the voxels within each LV, which involves randomly re-sampling participants with replacement and calculating the standard error of the saliences. Clusters larger than 20 contiguous voxels with a bootstrap ratio greater than three are reported (approximate P = .001, corrected at the whole brain level). Peaks within each cluster were specified based on the voxel with the highest bootstrap ratio in a 1 cm cube centred on the voxel. Given that PLS analysis identifies patterns of activity across the whole brain in a single step, corrections for multiple comparisons are not required.

We employed three types of PLS analysis in the current study. First, we took a data driven, mean-centered approach to examine the maximal differences between the six study conditions. Second, we conducted separate behavioral PLS analyses to identify patterns of brain activity associated with the online subjective ratings of vividness, emotional intensity, and perspective maintenance, as well as the reaction time to construct an AM or visualize the spatial location. For each behavioral PLS analysis, a cross-covariance matrix was computed across each trial including all participants and tasks by comparing a design matrix containing all voxels with a matrix containing the behavioral data in question.^1^ Third, we employed a seed PLS analyses to examine how visual perspective influenced the functional connectivity with the hippocampus over the time course of AM retrieval. We identified a seed region within the left anterior portion of hippocampus (MNI: *x* = −26, *y* = −6, *z* = - 22),from the spatiotemporal PLS analysis in time lag 6 (i.e. 15 s after cue onset), which was sensitive to changes in visual perspective during AM retrieval. The BOLD values in the hippocampal seed region for each participant at each time lag was then extracted and compared with activity in all other brain voxels separately within each condition.

## Results

### Behavioral Results

In general, we found that participants were slower to construct AMs than to visualize spatial locations, especially when adopting an atypical OE perspective (for *means* and *SD see* Table 1). There was a significant main effect of condition on the reaction time to construct an AM or to visualize the spatial location, *F* (2, 38) = 9.00, *p* = .001, η^2^_p_ = .32, reflecting slower reaction times in the OE (*M* = 4.30, *SD* = 1.65) and OB (*M* = 4.28, *SD* = 1.61) conditions compared to the spatial condition (*M* = 3.75, *SD* = 1.24), *p*′s < .05.

**Table 1.**
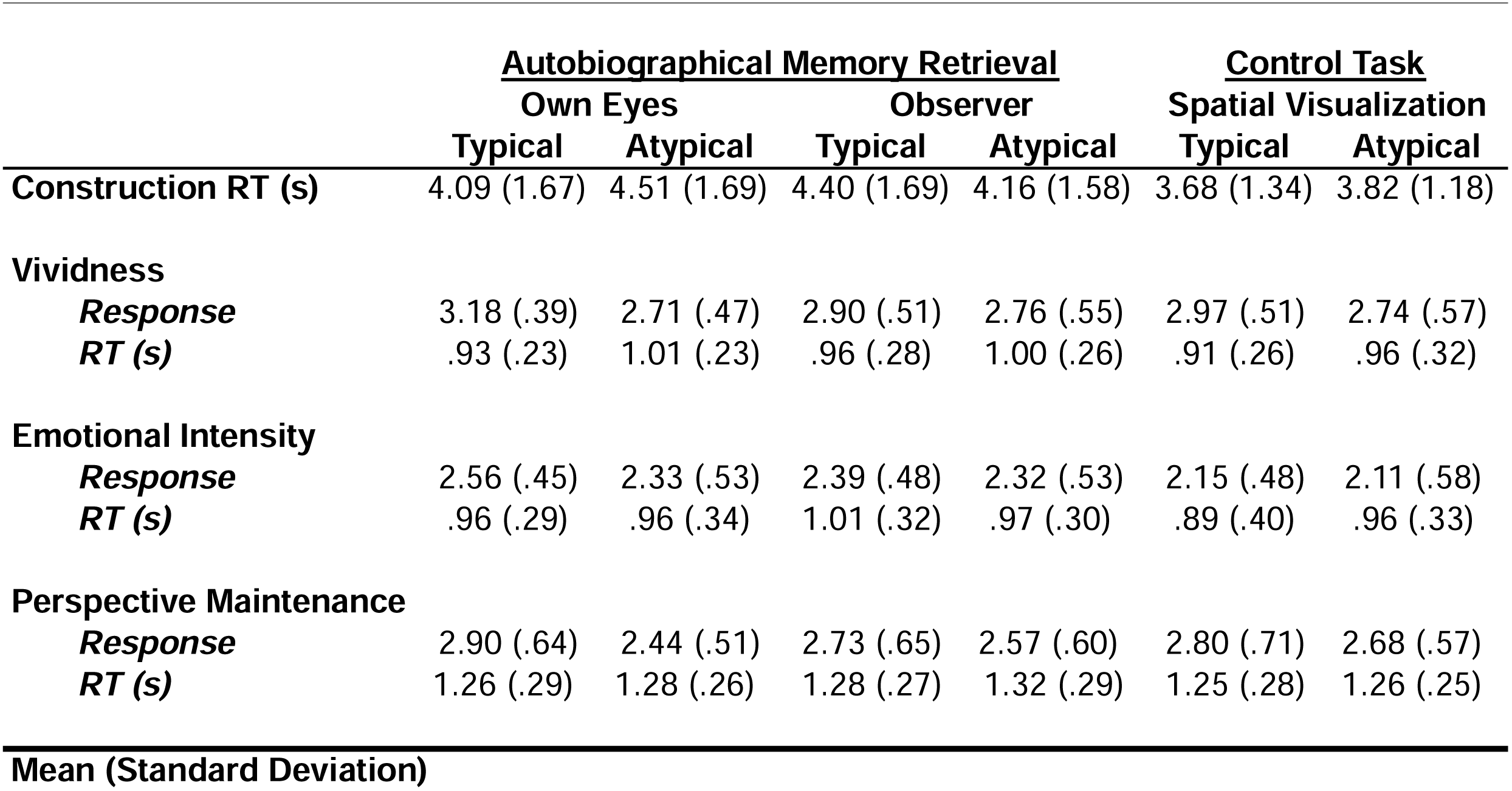
Behavioural Ratings and Reaction Time

The main effect was qualified by an interaction between condition and typicality, *F* (2, 36) = 8.75, *p* = .001, η^2^_p_ = .31. Follow-up tests indicated that the reaction time to construct events was slower in the atypical versus typical OE conditions, *p* = .01. For typical visual perspectives, reaction time was also slower in the OB compared to both the OE and Spatial conditions, *p*′s < .01. Whereas, for atypical visual perspectives, reaction time was slower for the OE compared to the Spatial Condition, *p* = .004.

Turning to the online subjective ratings, we found that typical versus atypical visual perspectives generally received higher subjective ratings (for *means* and *SDs see* Table 1). First, for vividness ratings there was a main effect of typicality, *F* (1, 18) = 32.34, *p* < .001, η^2^_p_ = .64, reflecting higher ratings for typical (M = 3.02, SD = .48) than atypical perspectives (*M* = 2.74, *SD* = .52). Reaction times for vividness ratings were also marginally faster for typical (*M* = .93, *SD* = .24) compared to atypical (*M* = .99, *SD* = .25) conditions, *F* (1,18) = 4.49, *p* = .05, η^2^_p_ = .20. The main effect of typicality was qualified by an interaction with condition, *F* (2, 36) = 4.21, *p* = .02, η^2^_p_ = .19. Follow-up tests indicated that vividness ratings were higher for typical compared to atypical perspectives in the OE and Spatial conditions, *p*’s < .05, but there was only a marginal effect in the OB condition, *p* = .06. Additionally, for typical perspectives, vividness ratings were higher in the OE compared to both the OB and Spatial conditions. However, there were no differences in vividness ratings between the conditions within atypical perspectives. Second, for emotional intensity ratings there was a main effect of typicality, *F* (1, 18) = 6.79, *p* = .02, η^2^_p_ = .27, with higher ratings for typical (*M* = 2.37, *SD* = .49) than atypical (*M* = 2.25, *SD* = .55) perspectives. There was also a main effect of condition, *F* (2, 36) = 15.82, p < .001, η^2^_p_ = .47, reflecting higher ratings in the OE (*M* = 2.44, *SD* = .49) and OB (*M* = 2.35, *SD* = .49) conditions compared to the Spatial conditions (M = 2.13, *SD* = .52), *p’*s < .005. Third, for perspective maintenance ratings there was a main effect of typicality, *F* (1, 18) = 7.32, *p* = .01, η^2^_p_ = .29, reflecting higher ratings for typical (*M* = 2.81, *SD* = .66) versus atypical (*M* = 2.56, *SD* = .56) perspectives. However, the main effect of typicality was qualified by an interaction with condition, *F* (2, 36) = 6.28, *p* = .005, η^2^_p_ = .26. Follow-up tests indicated higher perspective maintenance ratings in the typical versus atypical OE conditions, p = .004. Within typical perspectives, there were higher perspective maintenance ratings for OE compared to OB conditions, *p* = .03, whereas within atypical perspectives there were marginally lower ratings for OE than Spatial conditions, *p* = .05. There were no other main effects or interactions for subjective ratings or reaction times.

In sum, the behavioral findings suggest that the typicality of the perspective influenced the construction and online ratings in both the AM and Control tasks. In particular, AMs retrieved from typical OE perspectives were faster to construct and associated with greater vividness and ease of maintaining this perspective across the retrieval period. As expected, we found that emotional intensity was higher when retrieving AMs than visualizing a spatial location.

### Spatiotemporal PLS

The main goal of the study was to examine how visual perspective influences the construction and elaboration of AM retrieval. The spatiotemporal PLS analysis identified one significant LV, which accounted for 29.10% of the variance (*p* < .0001; see Table 2 and Figure 1). This LV maximally dissociated the typical OE and both OB conditions (negative brain scores) from the spatial and atypical OE conditions (positive brain scores). However, there were also significant differences in the weighting of the conditions on the LV. Specifically, the typical OE condition was assigned a greater negative brain score compared to both OB conditions, suggesting that the former contributed more to the overall pattern identified by the LV. Similarly, within the positive brain scores there was less loading on the atypical OE than the spatial control tasks, which also did not differ from one another.

**Table 2.**
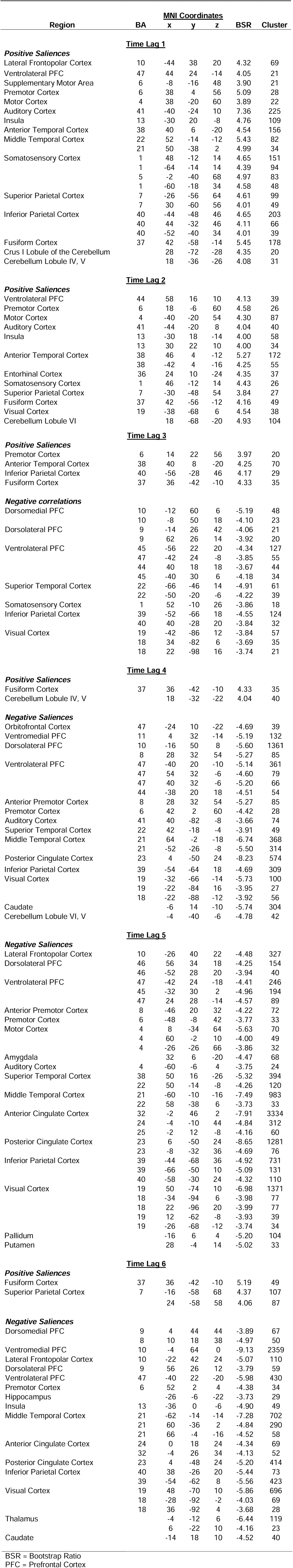
Spatiotemporal Task PLS

**Figure 1.**
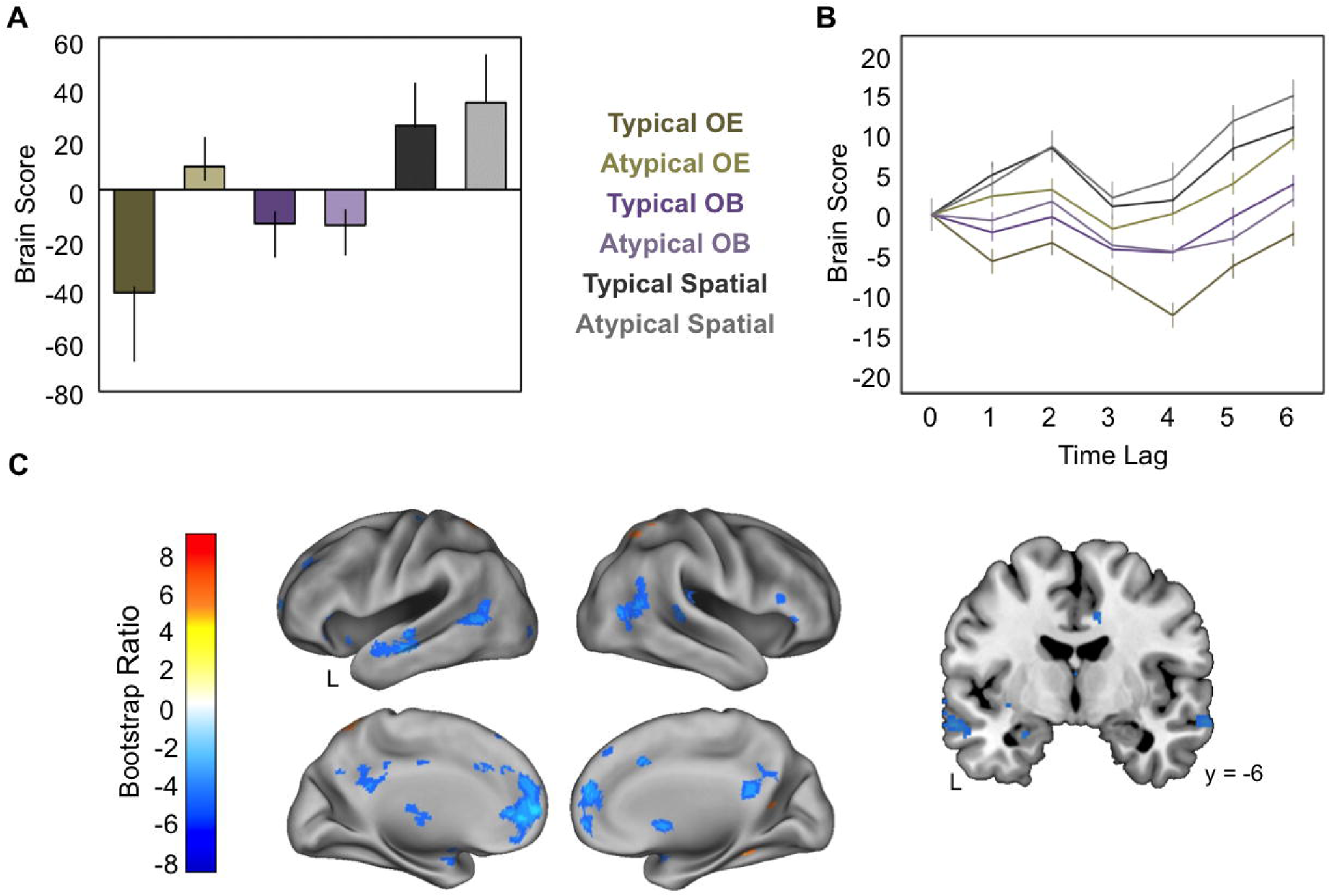
Spatiotemporal PLS Results. (A) The weighted average activation per condition across all voxels in all participants across the length of the retrieval period for the significant LV extracted from the spatiotemporal PLS. Error bars represent the 95% confidence interval and are based on bootstrap estimates. Positive brain scores are associated with spatial visualization and AM retrieval from an atypical OE perspective. Negative brain scores are associated with AM retrieval from typical OE and both OB perspectives. However, the typical OE perspective contributes more to negative brain scores compared to the OB conditions, which were not significantly different from one another. (B) The average brain score for each condition across each time lag within the trial. Each time lag corresponds to 2.5s (i.e., one TR). (C) Activation patterns corresponding to the positive and negative brain scores in time lag 6. All images depict a BSR threshold of +/- 3. OE = Own Eyes, OB = Observer.

Critically, the PLS approach also allowed us to distinguish how the LV differed during each time lag across the construction and elaboration phases. We found that the timing of the multivariate pattern associated with the positive and negative brain scores differed (see Figure 1B and Figure 1 in Supplementary material). Neural recruitment associated with the spatial and atypical OE conditions emerged earlier, peaking at time lag 2 (i.e., 5 s following cue onset). In contrast, neural recruitment associated with the typical OE and both OB conditions emerged later, peaking in response during time lag 4 (i.e., 9 s following cue onset) and persisting throughout the remainder of the trial. Given that the average length of construction was 4.11 s (*SD* = 1.46 s), these findings show that visual perspective influences the neural mechanisms that support later elaboration of AMs.

The spatial and atypical OE conditions recruited frontoparietal and lateral temporal cortices (see Table 2 and Figure 1C), which may reflect additional control processes required in these tasks relying on the retrieval of semantic information related to the AM or familiar location (for review see Svoboda and others 2006). In contrast, AMs retrieved in the typical OE and both OB conditions recruited anterior and posterior midline, MTL (including left anterior hippocampus and right amygdala), ventrolateral PFC, lateral temporal and inferior posterior parietal cortices (see Table 2 and Figure 1C), which overlap with regions frequently recruited during AM retrieval (for reviews see Cabeza and St Jacques 2007; Svoboda and others 2006). However, the pattern of neural recruitment contributed more to the typical OE condition compared to the OB conditions, demonstrating that these core AM regions are less engaged when elaborating upon memories from OB perspectives.

We conducted additional behavioral PLS analyses to determine whether the differences in the multivariate pattern during construction and elaboration overlapped with the pattern of neural activity sensitive to the reaction time to construct a memory or behavioral ratings (see Supplementary Tables 2 to 5). Separate behavioral PLS analyses were conducted on the reaction time to construct a memory or visualize the spatial location (see Supplementary Figure 2), as well as subjective ratings of perspective maintenance (see Supplementary Figure 3), vividness (see Supplementary Figure 4), and emotional intensity (see Supplementary Figure 5). These additional behavioral PLS analyses revealed that there was limited overlap with the spatiotemporal PLS (see Supplementary Figures 6 to 9), suggesting that the reported differences in visual perspective in the spatiotemporal PLS analysis did not overlap with regions sensitive to behavioral differences in construction reaction time, vividness, emotional intensity, or perspective maintenance.

### Hippocampal Seed PLS

We conducted a seed PLS analysis to examine whether functional connectivity with a left anterior hippocampus, which was recruited in both the typical OE and OB conditions, was sensitive to differences in visual perspective during construction and elaboration of AMs. Four significant LVs were found. LV1 (27.78% of variance, *p* =.001; see Supplementary Table 6 and Supplementary Figure 10) and LV4 (2.39% of variance, *p* =.01; see Supplementary Table 7 and Supplementary Figure 11) did not differentiate among the conditions. In contrast, LV2 (9.59% of variance, *p* = .03; Supplementary Figure 12A) distinguished atypical OB from typical OE conditions, and LV3 (10.93% of variance, *p* = .006; see Supplementary Figure 12B) distinguished the typical OE and atypical OB conditions. The temporal pattern in both LV2 and LV3 peaked at time lag 1 (i.e., 2.5 s following cue onset; during construction) and differences between the conditions persisted during elaboration across the remaining time lags (see Supplementary Figure 12C & 12D).

First, turning to differences between atypical OB and typical OE conditions shown in LV2 (see Table 3). During construction (i.e., time lag 1) there was greater hippocampal functional connectivity with left precuneus, angular gyrus, thalamus, right retrosplenial cortex, and bilateral posterior parahippocampal cortices (see Figure 2) . Notably, these regions belong to a posterior medial network (Ranganath and Ritchey 2012), which is linked to the translation of stored allocentric memory traces within the hippocampus to egocentric representations during long-term memory retrieval (Byrne and others 2007). In contrast, in the typical OE condition there was greater hippocampal functional connectivity with bilateral somatosensory cortices, perhaps reflecting greater access to interoceptive information when constructing memories from OE compared to OB perspectives (Eich and others 2009).

**Table 3.**
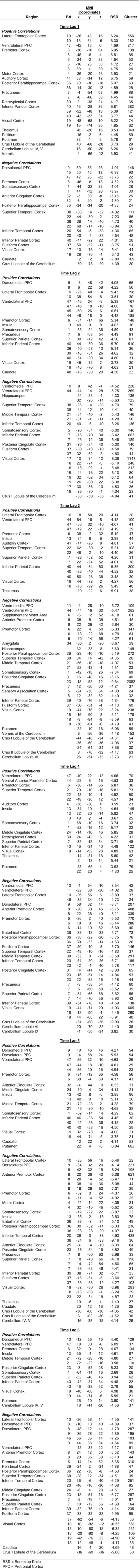
Hippocampus Seed PLS LV2

**Figure 2.**
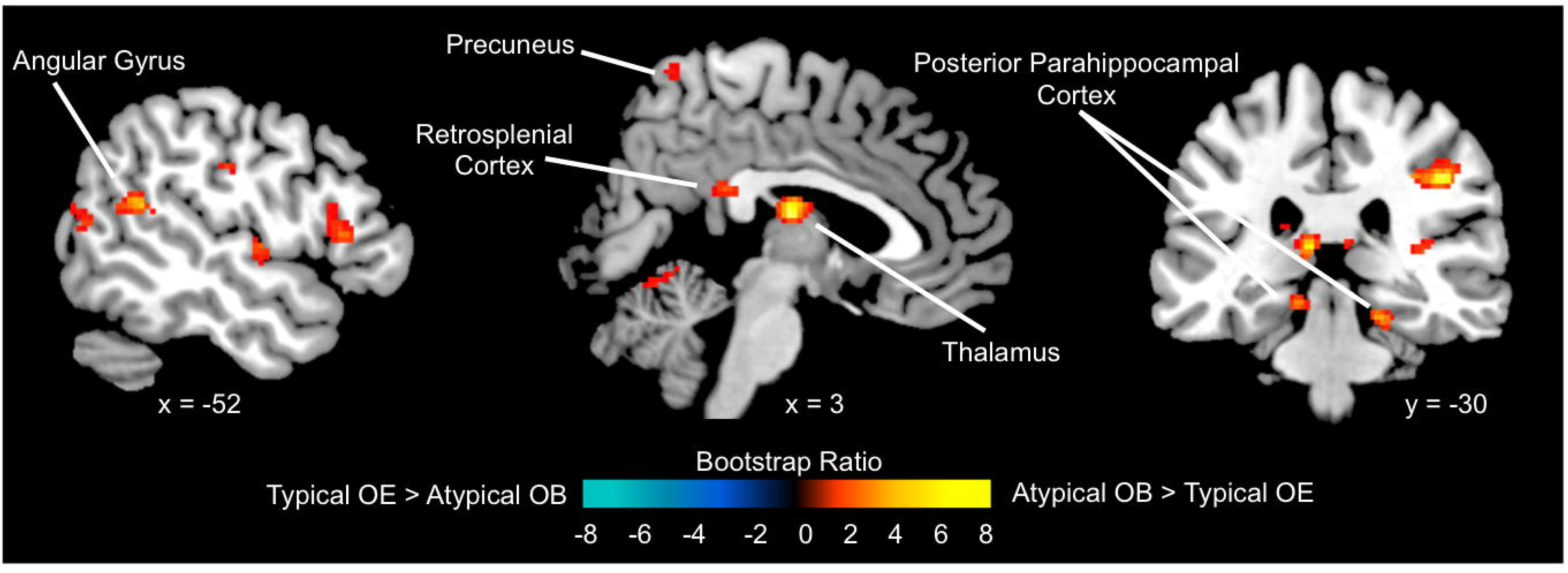
Hippocampus Functional Connectivity During Construction. The pattern of functional connectivity with the left anterior hippocampus identified in LV2 showing differences between the typical OE and atypical OB conditions during construction (i.e., time lag 1). OE = Own Eyes, OB = Observer. All images depict a BSR threshold of +/- 3.

In the ensuing elaboration period (i.e., time lags 2 to 6), for AMs retrieved in the typical OE compared to atypical OB perspectives, there was greater hippocampal functional connectivity within the MTL including right hippocampus, distinct sub-regions of the ipsilateral hippocampus, and posterior parahippocampal cortices, as well as left amygdala and entorhinal cortex (see Supplementary Figure 13). Hippocampal connectivity also extended outside of the MTL to ventromedial PFC and visual cortices, demonstrating greater integration among regions within a wider MTL network when elaborating upon AMs from typical OE perspectives (see Figure 3). In contrast, when elaborating upon AMs from atypical OB perspectives there was greater hippocampal functional connectivity with dorsomedial PFC. Additionally, as retrieval progressed from construction to elaboration functional connectivity between the hippocampus and precuneus became stronger in the typical OE compared to atypical OB conditions (i.e., time lags 3 to 6).

**Figure 3.**
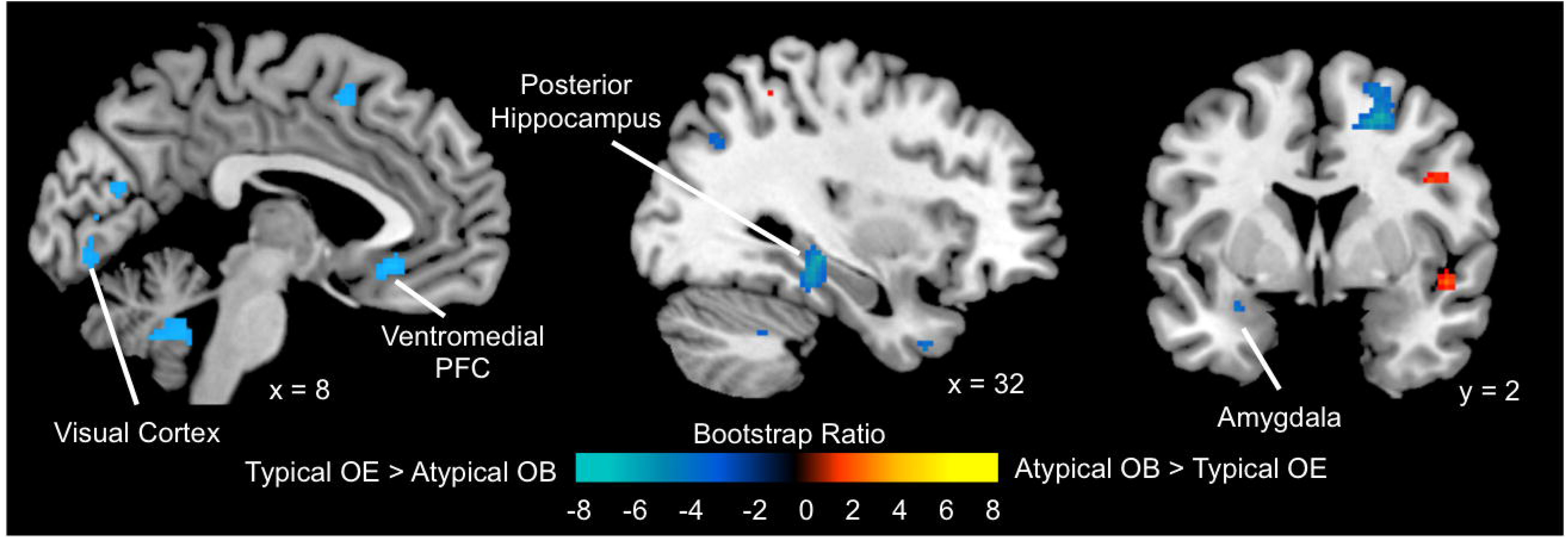
Hippocampus Functional Connectivity During Elaboration. The pattern of functional connectivity with the left anterior hippocampus identified in LV2 showing differences between typical OE and atypical OB conditions during elaboration (i.e., time lags 2 to 6; time lag 3 shown here). OE = Own Eyes, OB = Observer. All images depict a BSR threshold of +/- 3.

The seed PLS analysis also identified a third significant LV that reflected differences between both OB conditions and the typical spatial condition (see Table 4). During construction and early elaboration (i.e., time lags 1 to 2), the typical spatial condition was associated with greater functional connectivity between the hippocampus and a distributed set of cortical regions, including bilateral precuneus, prefrontal, and parietal cortices and right supplementary motor area (see Supplementary Figure 14). Substantial differences in hippocampal functional connectivity favoring the OB conditions did not manifest until time lag 3 and primarily implicated lateral temporal cortices. However, as elaboration progressed the OB conditions were characterized by greater within-MTL connectivity when compared to the spatial condition (see Supplementary Figure 15).

**Table 4.**
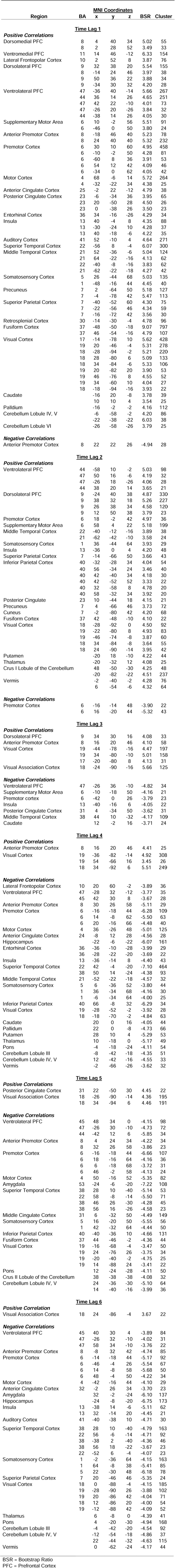
Hippocampus Seed PLS LV3

| Region | BA | MNI Coordinates |  |  | BSR | Cluster |
| --- | --- | --- | --- | --- | --- | --- |
|  |  | x | y | z |  |  |
| Time Lag 1 |  |  |  |  |  |  |
| Positive Correlations |  |  |  |  |  |  |
| Dorsomedial PFC | 8 | 4 | 40 | 34 | 5.02 | 55 |
|  | 8 | 2 | 28 | 52 | 3.49 | 33 |
| Ventromedial PFC | 11 | 14 | 46 | -12 | 6.33 | 154 |
| Lateral Frontopolar Cortex | 10 | 2 | 52 | 8 | 3.87 | 76 |
| Dorsolateral PFC | 9 | 32 | 38 | 20 | 5.54 | 155 |
|  | 8 | -14 | 24 | 46 | 3.97 | 38 |
|  | 9 | 50 | 36 | 22 | 3.88 | 34 |
|  | 9 | -34 | 30 | 32 | 4.20 | 28 |
| Ventrolateral PFC | 47 | -36 | 40 | -14 | 5.66 | 267 |
|  | 44 | 36 | 14 | 26 | 4.65 | 251 |
|  | 47 | 42 | 22 | -10 | 4.01 | 73 |
|  | 47 | -26 | 20 | -26 | 3.84 | 32 |
|  | 44 | -38 | 14 | 26 | 4.05 | 30 |
| Supplementary Motor Area | 6 | 10 | -2 | 56 | 5.51 | 91 |
|  | 6 | -46 | 0 | 50 | 3.80 | 24 |
| Anterior Premotor Cortex | 8 | -18 | 46 | 40 | 5.23 | 78 |
|  | 8 | 24 | 40 | 40 | 5.32 | 232 |
| Premotor Cortex | 6 | 30 | 10 | 60 | 4.95 | 458 |
|  | 6 | -10 | -2 | 50 | 4.28 | 81 |
|  | 6 | -60 | 8 | 36 | 3.91 | 53 |
|  | 6 | 54 | 12 | 42 | 4.09 | 46 |
|  | 6 | -34 | 0 | 62 | 4.05 | 42 |
| Motor Cortex | 4 | 68 | -6 | 14 | 5.72 | 264 |
|  | 4 | -32 | -22 | 34 | 4.38 | 25 |
| Anterior Cingulate Cortex | 25 | -2 | 22 | -12 | 4.79 | 38 |
| Posterior Cingulate Cortex | 23 | -6 | -34 | 36 | 3.95 | 65 |
|  | 23 | 20 | -50 | 28 | 4.50 | 26 |
|  | 23 | 0 | -38 | 26 | 3.50 | 23 |
| Entorhinal Cortex | 36 | 34 | -16 | -26 | 4.29 | 34 |
| Insula | 13 | 40 | -4 | 8 | 4.35 | 88 |
|  | 13 | -30 | -24 | 10 | 4.28 | 37 |
|  | 13 | 40 | -18 | -6 | 4.22 | 35 |
| Auditory Cortex | 41 | 52 | -10 | 4 | 4.64 | 271 |
| Superior Temporal Cortex | 22 | -56 | 8 | -4 | 6.07 | 300 |
| Middle Temporal Cortex | 21 | 52 | -36 | -6 | 5.04 | 124 |
|  | 21 | 64 | -22 | -16 | 4.13 | 62 |
|  | 22 | -40 | -8 | -16 | 3.83 | 62 |
|  | 21 | -62 | -22 | -18 | 4.27 | 42 |
| Somatosensory Cortex | 5 | 26 | -44 | 68 | 5.03 | 135 |
|  | 1 | -48 | -16 | 44 | 4.45 | 40 |
| Precuneus | 7 | 2 | -64 | 50 | 5.18 | 127 |
|  | 7 | -4 | -78 | 42 | 5.47 | 113 |
| Superior Parietal Cortex | 7 | -40 | -52 | 60 | 4.30 | 75 |
|  | 7 | -22 | -56 | 46 | 4.34 | 59 |
|  | 7 | -16 | -72 | 42 | 3.56 | 30 |
| Retrosplenial Cortex | 30 | -14 | -30 | -4 | 4.78 | 96 |
| Fusiform Cortex | 37 | -48 | -50 | -18 | 9.07 | 797 |
|  | 37 | 46 | -54 | -16 | 4.79 | 107 |
| Visual Cortex | 17 | -14 | -78 | 10 | 5.62 | 428 |
|  | 19 | 20 | -46 | -4 | 5.31 | 278 |
|  | 18 | -28 | -94 | -2 | 5.21 | 220 |
|  | 18 | 28 | -80 | 6 | 5.09 | 133 |
|  | 18 | 34 | -84 | -6 | 5.33 | 106 |
|  | 19 | 20 | -82 | 20 | 3.90 | 53 |
|  | 19 | 46 | -76 | 8 | 4.55 | 52 |
|  | 19 | 34 | -60 | 10 | 4.04 | 27 |
|  | 18 | -18 | -94 | -16 | 3.93 | 22 |
| Caudate |  | -16 | 20 | -8 | 3.78 | 39 |
|  |  | 10 | 10 | 4 | 3.54 | 25 |
| Palldium |  | -16 | -2 | -2 | 4.16 | 112 |
| Cerebellum Lobule IV, V |  | -6 | -58 | -2 | 4.20 | 86 |
|  |  | -22 | -38 | -22 | 6.03 | 38 |
| Cerebellum Lobule VI |  | -26 | -58 | -26 | 3.79 | 25 |
| **Negative Correlations** |  |  |  |  |  |  |
| Anterior Premotor Cortex | 8 | 22 | 22 | 26 | -4.94 | 28 |
| **Time Lag 2****Positive Correlations** |  |  |  |  |  |  |
| Ventrolateral PFC | 44 | -58 | 10 | -2 | 5.03 | 98 |
|  | 47 | 50 | 16 | -6 | 4.19 | 32 |
|  | 47 | -26 | 18 | -26 | 4.06 | 28 |
|  | 44 | 38 | 20 | 14 | 3.65 | 21 |
| Dorsolateral PFC | 9 | -24 | 40 | 38 | 4.87 | 330 |
|  | 9 | 38 | 32 | 18 | 5.26 | 227 |
|  | 9 | 26 | 38 | 34 | 4.58 | 120 |
|  | 9 | 12 | 50 | 38 | 3.79 | 23 |
| Premotor Cortex | 6 | 18 | -2 | 42 | 4.97 | 36 |
| Supplementary Motor Area | 6 | 58 | 4 | 22 | 5.18 | 199 |
| Middle Temporal Cortex | 22 | -40 | -12 | -16 | 3.89 | 38 |
|  | 21 | -62 | -42 | -10 | 3.58 | 24 |
| Somatosensory Cortex | 1 | 36 | -44 | 64 | 3.93 | 29 |
| Insula | 13 | -36 | 0 | 4 | 4.20 | 48 |
| Superior Parietal Cortex | 7 | -14 | -66 | 50 | 3.66 | 43 |
| Inferior Parietal Cortex | 40 | -32 | -28 | 34 | 4.04 | 54 |
|  | 40 | 56 | -34 | 24 | 3.46 | 40 |
|  | 40 | 42 | -40 | 34 | 4.18 | 30 |
|  | 40 | 42 | -52 | 52 | 3.33 | 22 |
|  | 40 | 32 | -58 | 8 | 4.78 | 20 |
|  | 40 | 58 | -32 | 34 | 3.92 | 20 |
| Posterior Cingulate | 23 | 10 | -44 | 18 | 4.15 | 21 |
| Precuneus | 7 | 4 | -66 | 46 | 3.73 | 72 |
| Cuneus | 7 | -2 | -80 | 42 | 4.20 | 68 |
| Fusiform Cortex | 37 | 42 | -48 | -10 | 4.10 | 22 |
| Visual Cortex | 18 | -28 | -92 | 0 | 4.50 | 92 |
|  | 19 | -22 | -80 | 8 | 4.93 | 83 |
|  | 19 | -46 | -74 | -8 | 3.87 | 60 |
|  | 18 | 34 | -84 | -8 | 3.64 | 55 |
|  | 18 | 24 | -90 | -14 | 3.95 | 42 |
| Putamen |  | -20 | 18 | -10 | 4.22 | 44 |
| Thalamus |  | -20 | -32 | 12 | 4.08 | 25 |
| Crus I Lobule of the Cerebellum |  | 48 | -50 | -30 | 4.25 | 48 |
|  |  | -20 | -82 | -22 | 4.51 | 237 |
| Vermis |  | -2 | -40 | -2 | 4.28 | 76 |
|  |  | 6 | -54 | -6 | 4.32 | 64 |
| **Negative Correlations** |  |  |  |  |  |  |
| Premotor Cortex | 6 | -16 | -14 | 48 | -3.90 | 22 |
|  | 6 | 16 | -20 | 44 | -5.32 | 43 |
| **Time Lag 3****Positive Correlations** |  |  |  |  |  |  |
| Dorsolateral PFC | 9 | 34 | 30 | 16 | 4.08 | 33 |
| Anterior Premotor Cortex | 8 | 16 | 20 | 46 | 4.10 | 58 |
| Visual Cortex | 19 | -44 | -78 | -16 | 4.47 | 197 |
|  | 19 | 34 | -80 | -10 | 5.01 | 158 |
|  | 17 | -20 | -80 | 8 | 4.13 | 31 |
| Visual Association Cortex | 18 | -24 | -90 | -16 | 5.66 | 125 |
| **Negative Correlations** |  |  |  |  |  |  |
| Ventrolateral PFC | 47 | -26 | 36 | -10 | -4.82 | 34 |
| Supplementary Motor Area | 6 | -10 | -18 | 50 | -4.16 | 21 |
| Premotor cortex | 6 | -42 | 0 | 26 | -3.79 | 23 |
| Insula | 13 | -40 | -16 | 6 | -4.05 | 22 |
| Posterior Cingulate Cortex | 31 | 4 | -34 | 50 | -3.62 | 31 |
| Middle Temporal Cortex | 38 | 44 | 10 | -32 | -4.17 | 109 |
| Caudate |  | 12 | -2 | 16 | -3.71 | 24 |
| **Time Lag 4****Positive Correlations** |  |  |  |  |  |  |
| Anterior Premotor Cortex | 8 | 16 | 20 | 46 | 4.41 | 25 |
| Visual Cortex | 19 | -36 | -82 | -14 | 4.92 | 308 |
|  | 19 | 54 | -66 | 16 | 3.45 | 26 |
|  | 18 | 34 | -92 | 6 | 5.51 | 249 |
| **Negative Correlations** |  |  |  |  |  |  |
| Lateral Frontopolar Cortex | 10 | 20 | 60 | -2 | -3.89 | 36 |
| Ventrolateral PFC | 47 | -28 | 32 | -12 | -3.77 | 35 |
|  | 45 | 42 | 30 | 8 | -3.67 | 28 |
| Anterior Premotor Cortex | 8 | 30 | 26 | 58 | -5.11 | 29 |
| Premotor Cortex | 6 | -16 | -18 | 44 | -6.28 | 109 |
|  | 6 | 14 | -8 | 62 | -5.50 | 63 |
|  | 6 | 20 | -16 | 66 | -4.48 | 40 |
| Motor Cortex | 4 | 36 | -26 | 48 | -5.01 | 125 |
| Anterior Cingulate Cortex | 24 | -8 | 12 | 28 | -4.56 | 28 |
| Hippocampus |  | -22 | -6 | -22 | -6.07 | 161 |
| Entorhinal Cortex | 36 | -36 | -10 | -28 | -3.99 | 29 |
|  | 36 | -28 | -22 | -20 | -3.69 | 22 |
| Insula | 13 | -36 | -14 | 8 | -4.40 | 43 |
| Superior Temporal Cortex | 22 | 42 | -4 | -20 | -7.10 | 464 |
|  | 38 | 50 | 14 | -24 | -4.38 | 93 |
| Middle Temporal Cortex | 21 | -52 | -22 | -18 | -4.57 | 32 |
| Somatosensory Cortex | 5 | 6 | -36 | 52 | -3.80 | 44 |
|  | 1 | 36 | -34 | 68 | -4.16 | 30 |
|  | 1 | -6 | -34 | 64 | -4.00 | 25 |
| Inferior Parietal Cortex | 40 | 66 | -8 | 32 | -6.29 | 34 |
| Visual Cortex | 19 | -28 | -52 | -2 | -3.92 | 28 |
|  | 18 | -18 | -70 | -2 | -4.84 | 63 |
| Caudate |  | 20 | 0 | 16 | -4.05 | 44 |
| Pallidum |  | 22 | 0 | -8 | -4.73 | 66 |
| Putamen |  | 28 | 10 | 4 | -5.29 | 53 |
| Thalamus |  | 10 | -18 | 0 | -5.17 | 49 |
| Pons |  | -4 | -18 | -24 | -4.11 | 54 |
| Cerebellum Lobule III |  | -8 | -42 | -18 | -4.35 | 51 |
| Cerebellum Lobule IV, V |  | 12 | -42 | -16 | -4.55 | 33 |
| Vermis |  | -2 | -66 | -26 | -3.62 | 32 |
| **Time Lag 5****Positive Correlations** |  |  |  |  |  |  |
| Posterior Cingulate Cortex | 31 | 22 | -50 | 30 | 4.45 | 22 |
| Visual Association Cortex | 18 | -26 | -90 | -14 | 4.36 | 195 |
|  | 18 | 34 | -94 | 6 | 4.46 | 191 |
| **Negative Correlations** |  |  |  |  |  |  |
| Ventrolateral PFC | 45 | 48 | 34 | 0 | -4.15 | 98 |
|  | 47 | -26 | 30 | -10 | -4.73 | 72 |
|  | 44 | -42 | 12 | 6 | -5.85 | 34 |
| Anterior Premotor Cortex | 8 | 4 | 24 | 34 | -4.22 | 34 |
|  | 8 | 32 | 26 | 58 | -3.86 | 23 |
| Premotor Cortex | 6 | -16 | -18 | 44 | -6.66 | 107 |
|  | 6 | 18 | -16 | 64 | -4.16 | 36 |
|  | 6 | 6 | -18 | 68 | -3.72 | 31 |
|  | 6 | 46 | -2 | 58 | -4.13 | 24 |
| Motor Cortex | 4 | 50 | -16 | 52 | -5.35 | 82 |
| Amygdala | 53 | -24 | -6 | -20 | -7.22 | 108 |
| Superior Temporal Cortex | 38 | 28 | 10 | -40 | -5.14 | 53 |
|  | 22 | 58 | -8 | -14 | -5.50 | 71 |
|  | 38 | 46 | 26 | -30 | -4.28 | 45 |
|  | 38 | 56 | 16 | -26 | -4.58 | 23 |
| Middle Cingulate Cortex | 31 | 6 | -32 | 50 | -4.49 | 149 |
| Somatosensory Cortex | 5 | 16 | -20 | 50 | -5.55 | 56 |
|  | 1 | 42 | -32 | 64 | -4.44 | 50 |
| Inferior Parietal Cortex | 40 | -40 | -36 | 10 | -4.66 | 131 |
| Fusiform Cortex | 37 | 44 | -46 | -2 | -4.36 | 44 |
| Visual Cortex | 19 | -16 | -68 | -4 | -3.47 | 50 |
|  | 19 | 24 | -76 | 26 | -3.75 | 34 |
|  | 19 | -20 | -40 | -2 | -4.75 | 25 |
|  | 19 | 14 | -88 | 24 | -3.41 | 22 |
| Pons |  | 12 | -24 | -28 | -4.11 | 50 |
| Crus II Lobule of the Cerebellum |  | 38 | -38 | -38 | -4.08 | 32 |
| Cerebellum Lobule IV, V |  | 24 | -36 | -30 | -5.10 | 64 |
|  |  | 14 | -40 | -16 | -3.99 | 36 |
| **Time Lag 6*****Positive Correlation*** |  |  |  |  |  |  |
| Visual Association Cortex | 18 | 24 | -86 | -4 | 3.67 | 22 |
| ***Negative Correlations*** |  |  |  |  |  |  |
| Ventrolateral PFC | 45 | 40 | 30 | 4 | -3.89 | 84 |
|  | 47 | -26 | 32 | -10 | -4.02 | 31 |
|  | 47 | 58 | 34 | -10 | -3.76 | 22 |
| Anterior Premotor Cortex | 8 | -8 | 32 | 42 | -4.74 | 85 |
| Premotor Cortex | 6 | -14 | -18 | 44 | -5.17 | 92 |
|  | 6 | -46 | -4 | 26 | -5.54 | 67 |
|  | 6 | 14 | -8 | 58 | -5.68 | 50 |
|  | 6 | 48 | -4 | 50 | -4.22 | 34 |
| Motor Cortex | 4 | -42 | -16 | 44 | -4.10 | 29 |
| Anterior Cingulate Cortex | 32 | -2 | 26 | 34 | -3.70 | 23 |
| Amygdala |  | 32 | -2 | -24 | -6.10 | 137 |
| Hippocampus |  | -24 | -8 | -20 | -6.75 | 173 |
| Insula | 13 | -38 | 14 | -6 | -5.11 | 62 |
|  | 13 | 32 | -14 | 20 | -4.45 | 21 |
| Auditory Cortex | 41 | -40 | -38 | 10 | -4.71 | 30 |
| Superior Temporal Cortex | 38 | 28 | 10 | -40 | -4.79 | 163 |
|  | 22 | 56 | -6 | -14 | -4.71 | 92 |
|  | 38 | -38 | 2 | -40 | -4.36 | 46 |
|  | 38 | 56 | 18 | -22 | -3.67 | 23 |
|  | 22 | -52 | 6 | -4 | -4.07 | 23 |
| Somatosensory Cortex | 1 | -2 | -36 | 64 | -4.15 | 163 |
|  | 1 | 64 | -8 | 38 | -5.41 | 85 |
|  | 5 | 22 | -30 | 48 | -6.18 | 78 |
| Superior Parietal Cortex | 7 | 20 | -46 | 46 | -5.35 | 24 |
| Visual Cortex | 18 | 0 | -88 | -4 | -4.15 | 185 |
|  | 19 | -28 | -90 | 26 | -3.88 | 102 |
|  | 19 | 20 | -86 | 42 | -4.04 | 71 |
|  | 18 | 12 | -86 | 20 | -4.00 | 54 |
|  | 19 | -12 | -88 | 42 | -4.09 | 52 |
| Thalamus |  | 6 | -8 | 0 | -4.39 | 41 |
| Pons |  | 4 | -20 | -30 | -4.94 | 168 |
| Cerebellum Lobule III |  | -4 | -42 | -20 | -4.54 | 92 |
| Cerebellum Lobule IV, V |  | -12 | -54 | -18 | -4.86 | 37 |
|  |  | 22 | -44 | -32 | -4.63 | 115 |
| Vermis |  | 0 | -62 | -24 | -4.17 | 44 |

BSR = Bootstrap Ratio
PFC = Prefrontal Cortex

In sum, the findings from the seed PLS analysis demonstrate that functional connectivity with the left anterior hippocampus is altered during construction and elaboration phases when AMs are retrieved from atypical OB relative to typical OE perspectives, and when comparing AM retrieval from an OB perspective relative to spatial visualization of the proximal aspects of a familiar location. The construction of AMs from atypical OB perspectives involved greater hippocampal functional connectivity with a posterior medial network (i.e., thalamus, retrosplenial cortex, precuneus, and angular gyrus), whereas elaboration of AMs from typical OE perspectives involved hippocampal connectivity with an MTL network (i.e., within-MTL and ventromedial PFC). Further differences between the OB and typical spatial conditions demonstrated a continuum of within-MTL connectivity whereby connectivity during elaboration was strongest in the typical OE condition, moderate in the OB conditions, and weakest in the typical spatial condition.

## Discussion

Our findings reveal that visual perspective shapes the spatiotemporal dynamics of the brain networks that underlie AM retrieval. Theories of memory suggest that the initial retrieval of events involves a constructive process of searching for, accessing, and assembling stored information, which is followed by the re-experience and elaboration of the sensory and perceptual qualities of memories (Cabeza and St Jacques 2007; Conway and Pleydell-Pearce 2000). Using a multivariate analytical technique, our results provide evidence that the particular visual perspective adopted during retrieval influences how memories are constructed and elaborated. We found that adopting a particular visual perspective affected how memories were elaborated upon, as reflected by increased neural recruitment of a core AM retrieval network (i.e., MTL, anterior and posterior midline regions, lateral PFC and posterior parietal cortices) for OE compared to OB perspectives. However, hippocampal functional connectivity analyses also revealed key differences in how visual perspective influenced the integration among neural regions during both construction and elaboration of AMs. There was stronger hippocampal connectivity with a posterior medial network during construction of AMs from OB perspectives, but stronger connectivity with an MTL network during elaboration of AMs from OE perspectives. Taken together, these results demonstrate that visual perspective affects how and when different neocortical systems guide memory retrieval.

Only a handful of functional neuroimaging studies have investigated how visual perspective influences long-term memory retrieval (Eich and others 2009; Grol and others 2017; St Jacques and others 2018; St Jacques and others 2017). Eich and colleagues (2009) found greater recruitment of the amygdala for OE perspectives coupled with reduced neural recruitment of somato-motor and insular cortices for OB perspectives, which they suggested reflected reductions in emotion and embodiment when adopting an OB perspective during retrieval. In contrast, Grol and colleagues (2017) found greater recruitment of precuneus and temporoparietal junction for OB compared to OE perspectives, which they linked to increased involvement of self-referential and visuospatial processing for OB perspectives. In the current study we did not find any additional regions, outside of a common AM retrieval network, that contributed more for OE compared to OB perspectives. However, there are several methodological differences between the current study and previous research that could explain these different results. For example, Eich and colleagues (2009) used complex lab-based events based on physical actions that may have depended more on somato-motor and insular cortices when compared with the AMs used in the present study, which varied in content and type of event retrieved. Additionally, here we directly elicited AMs associated with OE and OB perspectives, which may have reduced demands to shift to an alternative visual perspective that are supported by the precuneus (e.g., St Jacques and others 2017). Instead, our findings suggest that visual perspective influences how particular brain regions interact with the hippocampus across both construction and elaboration phases of AM retrieval, contributing to later changes in neural recruitment in a core memory retrieval network.

Visual perspective influenced hippocampal functional connectivity during both the construction and elaboration phases of AM retrieval. The hippocampus is crucial for binding together disparate elements of memories that support mental constructions and contribute to vivid recall (for recent reviews see Maguire and others 2016; Robin 2018; Sheldon and Levine 2016). During memory retrieval the hippocampus acts as a hub to coordinate the spatiotemporal dynamics of construction and elaboration (McCormick and others 2015) and the timing of multiple large-scale brain networks (Inman and others 2018; St Jacques and others 2013). Additionally, previous research has demonstrated that the hippocampus is recruited during memory retrieval from both OE and OB perspectives (Eich and others 2009; Grol and others 2017; St Jacques and others 2018; St Jacques and others 2017), but is altered when memories are formed from OB perspectives (Bergouignan and others 2014). Here, we focused on a seed region placed in the anterior portion of the hippocampus that was engaged during AM retrieval from both OE and OB perspectives. Recent theories have postulated functional specialization along the long axis of the hippocampus, with anterior portions supporting the flexible construction of mental scenarios and more posterior portions supporting the detailed retrieval of local aspects of an event (for reviews see Addis and Schacter 2012; Moscovitch and others 2016; Poppenk and others 2013; Schlichting and Preston 2015; Zeidman and Maguire 2016). Our findings demonstrate that OE and OB perspectives differentially affected the pattern of co-activation with the anterior hippocampus, suggesting that an important determinant of the transient connectivity supporting AM (e.g., McCormick and others 2015) is the particular visual perspective adopted during retrieval.

During the initial construction of AMs from OB compared to OE perspectives there was greater integration between the hippocampus and a posterior medial network that included thalamus, posterior parahippocampal cortex, retrosplenial cortex, precuneus and angular gyrus. The posterior medial network is thought to support the construction of situational models of events by assembling spatial and temporal contextual information from a particular egocentric perspective (Ranganath and Ritchey 2012), which contributes to the recollection of memories as well as related processes of scene construction and imagination of hypothetical events (Robin 2018). Computational theories of memory and imagery additionally specify that interactions between the hippocampus and other regions within the posterior medial network enable stored allocentric memory representations to be transformed to egocentric ones during long-term memory retrieval (Byrne and others 2007). Our findings reveal that the particular egocentric perspective adopted when constructing events modulates how and when the posterior medial network interacts with the hippocampus. Atypical (i.e., floor-level) but not typical (i.e., eye-level) OB perspectives were supported by greater hippocampal integration with the posterior medial network when compared to typical OE perspectives. One reason may be that adopting an atypical OB perspective required greater translation between allocentric and egocentric representations in memory and placed greater demands on the transformation circuit (e.g., Dhindsa and others 2014; Lambrey and others 2012), consistent with evidence that actively shifting from a dominant to an alternative visual perspective during AM retrieval involves greater recruitment of precuneus and angular gyrus (St Jacques and others 2017). These findings highlight the need to better understand the variety of visual perspectives that can be taken during memory retrieval (Rice and Rubin 2011), in line with research demonstrating that multiple visual perspectives can be flexibly adopted during conscious experience and impact how memories are formed and later retrieved (Bergouignan and others 2014).

As construction moved to elaboration there was a reversal in the pattern of hippocampal-precuneus functional connectivity favouring OE compared to OB perspectives. The precuneus is associated with visual imagery processes during AM retrieval (Ahmed and others 2018; Fuentemilla and others 2014) that are recruited during elaboration (Daselaar and others 2008), in line with theories of AM that postulate that perceptual information contributing to remembering occurs later during retrieval (Conway and Pleydell-Pearce 2000). Moreover, vivid retrieval of AMs has also been shown to be associated with higher phase-synchrony in theta-oscillations between the precuneus and hippocampus (Fuentemilla and others 2014). Our findings show that the nature and timing of integration between the hippocampus and precuneus depends upon the particular visual perspective adopted during retrieval. Recent evidence suggests that the precuneus supports a more general ability to mentally orientate the self within a large-scale environment (Peer and others 2015), consistent with the involvement of this region in multiple domains (Cavanna and Trimble 2006). Thus, one interpretation of the current findings is that OB perspectives during AM retrieval place early demands on mental orientation because adopting this non-dominant perspective likely requires restructuring visual imagery from a novel egocentric vantage point, whereas OE perspective may draw upon mental orientation later as perceptual information is retrieved during elaboration.

During elaboration, AM retrieval from OE compared to OB perspectives was also supported by greater integration within an MTL network that included ventromedial PFC, amygdala, posterior parahippocampal, hippocampal and entorhinal cortices, as well as visual cortices. Additionally, a similar pattern of within-MTL connectivity also contributed more to AMs retrieved from OB perspectives when compared to spatial visualization of proximal aspects of a familiar location, suggesting greater integration among these regions during retrieval of specific events. Similarly, St. Jacques and colleagues (2013) showed greater integration in an MTL network centered on the hippocampus among people who spontaneously recalled more AMs from stronger OE perspectives. Here we replicate these findings but also significantly extend them by demonstrating that people can flexibly engage this network in the service of retrieving memories from a specific perspective. The MTL network is implicated in the retrieval of episodic information contributing to the recollection or visualization of mental scenes and hypothetical events (Andrews-Hanna and others 2010; Kahn and others 2008). In particular, amygdalar-hippocampal interactions are thought to contribute to the recollection of memories based on salient item-specific details, which may be further supported by the recapitulation of perceptual information in the ventral visual stream via projections to the hippocampus from the entorhinal cortex (for review see Buchanan 2007; Phelps and Sharot 2008; Yonelinas and Ritchey 2015). The stronger co-activation among these regions when retrieving AMs from OE perspectives likely contributed to higher subjective ratings of vividness when compared to OB perspectives. We also found co-activation between the anterior hippocampus and ventromedial PFC, a region linked to conceptual aspects of self-reference and affective value (e.g., Bergstrom and others 2015; Lin and others 2016) that enable the formation of abstract mental models or schemas about the world and oneself in order to extract meaning to guide behavior (for reviews see D’Argembeau 2013; Gilboa and Marlatte 2017; Morton and others 2017; Roy and others 2012). During retrieval, interactions between the ventromedial PFC and the anterior hippocampus are thought to contribute to updating of reactivated memories guided by abstract memory representations (Schlichting and Preston 2015), which may contribute to the transformation of memories overtime (Moscovitch and others 2016). In contrast, elaboration of AMs from OB perspectives was associated with increased hippocampal connectivity with dorsomedial PFC, which is linked to the processing of social information related to other people (for metaanalysis see Denny and others 2012). For example, St. Jacques and colleagues (2011a) found that dorsomedial PFC is recruited to a greater extent when people were asked to understand another person’s perspective, whereas ventromedial PFC was recruited more during AM retrieval for events cued from an own eyes perspective (also see Rabin and others 2010). The ventral versus dorsal distinction in the medial PFC found here could reflect differences in how AM retrieval from OE and OB perspectives are guided by self-related schemas (e.g., Libby and Eibach 2011; Sutin and Robins 2008). The ability to adopt multiple egocentric perspectives that vary in their self-distance in memories offers a potential bridge between self- and other-related representations that enable us to understand mental states in others.

Visual perspective also influenced neural recruitment in a core AM retrieval network during elaboration in a similar way to other aspects of mental imagery that tend to occur later during retrieval (Daselaar and others 2008), and perhaps due to changes in the functional integration with the hippocampus (McCormick and others 2015). Critically, the pattern of neural recruitment during elaboration of AMs from multiple visual perspectives was distinguished from general differences in visualizing a scene from alternative viewpoints. Moreover, we also showed that this pattern was not directly related to behavioural differences in the subjective experience associated with retrieval, because there was little overlap between neural regions that contributed to behavior and those that distinguished OE and OB perspectives. Behavioural studies have demonstrated that visual perspective alters the phenomenology of memories during retrieval (e.g., Berntsen and Rubin 2006) as also shown here, which is thought to contribute to differences in neural recruitment as has sometimes been reported (Eich and others 2009; Grol and others 2017). In the current study, the slower construction times for AMs experienced from OB perspectives also shortened the subsequent elaboration period and may have obscured the association between visual perspective and subjective aspects of elaboration (i.e., vividness and emotional intensity). Instead, our findings point to the differences in hippocampal interactions with the posterior medial network and a wider MTL network, which could have contributed to subjective changes in AM retrieval due to visual perspective.

## Conclusion

Egocentric perspective is a defining feature of memories for events (Byrne and others 2007; Robin 2018; Rubin and Umanath 2015), but this self-centered aspect of remembering has been elusive to investigation due to its ubiquitous nature (Prebble and others 2013). Here, by manipulating multiple visual perspectives during AM retrieval, we demonstrate for the first time how egocentric perspective alters the neural mechanisms that contribute to the time-course of AM retrieval. We found that OE and OB perspectives were associated with identical patterns of activation in the AM retrieval network during elaboration, but to a lesser extent for OB perspectives. However, functional connectivity with the hippocampus revealed earlier posterior medial network involvement when adopting an OB perspective, coupled with greater engagement of an MTL network when adopting an OE perspective during elaboration. The current findings contribute to research on how visual perspective shapes memories during retrieval (Marcotti and St Jacques 2018; Sekiguchi and Nonaka 2014; St Jacques and others 2017), in line with theories of memory that propose retrieval is an active process that can reshape memories (e.g., Moscovitch and others 2016; Schacter and others 2011; Scully and others 2017). Better understanding of the neural mechanisms that support the fundamental capacity to understand ourselves from multiple perspectives when remembering the past could also contribute to research on how we flexibly understand the perspective of others (Carrington and Bailey 2009).

## Acknowledgments

We thank Petra Marcotti and Jazel Keir for their assistance with participant testing, and radiographers Janice Bush, James Hunter, and Marta Andrade, at the Clinical Imaging Sciences Centre, University of Sussex, for their assistance with fMRI data collection.

## Footnotes

1 Data from one participant was excluded in the behavioral PLS analyses on subjective ratings, due to an insufficient number of behavioral responses.

